# Temporal prediction elicits rhythmic pre-activation of relevant sensory cortices

**DOI:** 10.1101/2020.10.20.347005

**Authors:** Louise Catheryne Barne, André Mascioli Cravo, Floris P. de Lange, Eelke Spaak

## Abstract

Being able to anticipate events before they happen facilitates stimulus processing. The anticipation of the contents of events is thought to be implemented by the elicitation of prestimulus templates in sensory cortex. In contrast, the anticipation of the timing of events is typically associated with entrainment of neural oscillations. It is so far unknown whether and in which conditions temporal expectations interact with feature-based expectations, and, consequently, whether entrainment modulates the generation of content-specific sensory templates. In this study, we investigated the role of temporal expectations in a sensory discrimination task. We presented participants with rhythmically interleaved visual and auditory streams of relevant and irrelevant stimuli while measuring neural activity using magnetoencephalography. We found no evidence that rhythmic stimulation induced prestimulus feature templates. However, we did observe clear anticipatory rhythmic pre-activation of the relevant sensory cortices. This oscillatory activity peaked at behaviourally relevant, in-phase, intervals. Our results suggest that temporal expectations about stimulus features do not behave similarly to explicitly cued, non-rhythmic, expectations; yet elicit a distinct form of modality-specific pre-activation.

**Graphical abstract:** 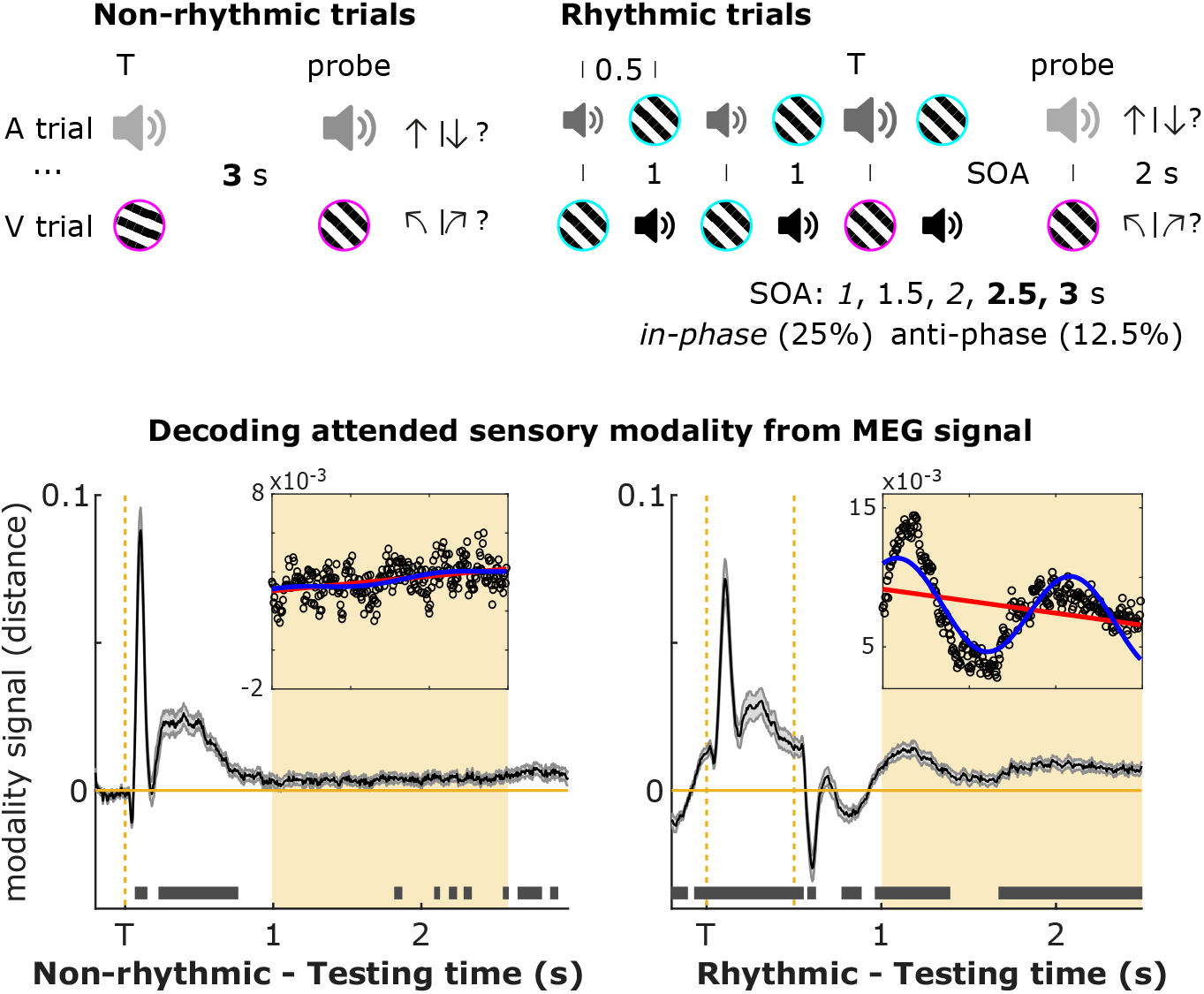

The brain extracts temporal regularities from the environment to anticipate upcoming events. Furthermore, with prior knowledge about their contents, the brain is thought to leverage this by instantiating anticipatory sensory templates. We investigated if sensory templates occur in response to a rhythmic stimulus stream with predictable temporal structure. We found that temporal rhythmic predictions did not induce sensory templates, but rather modulated the excitability in early sensory cortices.

## Introduction

Predicting upcoming events enables efficient resource allocation and can lead to behavioural benefits and neural processing improvements (Summerfield & De Lange, 2014; de Lange, Heilbron, & Kok, 2018). These predictions, or expectations, can come from various sources. For example, predictions can be the result of an explicit instruction (“when you see X, expect Y”), they can be (implicitly) inferred from the statistics of the world (Oliva & Torralba, 2007; Bar, 2004; Seriès & Seitz, 2013; Spaak & de Lange, 2020), or they can stem from temporal regularities in the sensory input (de Lange et al., 2018; Nobre & Van Ede, 2018). One proposed mechanism of how expectations can modulate perception is by inducing sensory templates through prestimulus baseline increases in sensory neurons tuned to the features of expected stimuli (SanMiguel, Widmann, Bendixen, Trujillo-Barreto, & Schröger, 2013; Kok, Failing, & de Lange, 2014; Kok, Mostert, & De Lange, 2017). A recent study using multivariate decoding techniques in MEG signal showed that an auditory cue that allowed observers to form an expectation of a particular grating orientation induced a visual prestimulus activation similar to the feature-specific response evoked by the actual visual stimulation (Kok et al., 2017).

It is unknown whether a similar mechanism is at play in anticipating the likely *time* of relevant events. Several studies have found faster and more accurate responses when stimuli are expected in time (Nobre, Correa, & Coull, 2007; Rohenkohl, Cravo, Wyart, & Nobre, 2012; Nobre, 2001). Studies in human and non-human primates have shown that neural populations in primary cortical regions can synchronise in frequency and phase to external rhythmic temporal patterns (Lakatos, Karmos, Mehta, Ulbert, & Schroeder, 2008; Schroeder & Lakatos, 2009; Lakatos et al., 2013; Besle et al., 2011; Cravo, Rohenkohl, Wyart, & Nobre, 2013; Henry, Herrmann, & Obleser, 2014). High and low neuronal ensemble excitability states could be entrained to stimulus timing in such a way that optimal phases of processing become aligned with the expected moments of task-relevant stimuli (Lakatos et al., 2008; Schroeder & Lakatos, 2009; Lakatos et al., 2013).

Generally, entrainment is marked by a strong phase coherence of neural signals at the stimulated frequency and by correlations between phase and attention and/or behavioural performance. However, there is no consensus on the definition of neural oscillatory entrainment (Obleser, Henry, & Lakatos, 2017; Breska & Deouell, 2017; Lakatos, Gross, & Thut, 2019; Haegens, 2020). Critically, most previous studies have used a stimulus-driven paradigm (i.e., testing entrainment at the same time when driving stimuli are present), which makes conclusions about the underlying mechanism hard to interpret, especially in non-invasive human studies (Haegens & Golumbic, 2018). There is an increasing debate whether the oscillatory modulation is purely due to superimposed evoked responses (Capilla, Pazo-Alvarez, Darriba, Campo, & Gross, 2011; van Diepen & Mazaheri, 2018) or to true endogenous oscillatory entrainment (Doelling, Assaneo, Bevilacqua, Pesaran, & Poeppel, 2019). A few studies have reported behavioural and neural oscillatory modulations persisting after the offset of rhythmic stimulation (Lakatos et al., 2013; Spaak, de Lange, & Jensen, 2014), thus providing stronger evidence for the importance of neural entrainment.

The existence of these two mechanisms for preparing for upcoming stimuli (prestimulus templates in response to explicit cues, neural entrainment in response to rhythmically induced temporal expectations) raises the interesting question of whether and how these two mechanisms interact or complement each other. We here aim to shed light on this question. Specifically, we hypothesized that stimulus-specific sensory templates might emerge at the relevant phases of the entraining signal, i.e., the time points of expected stimulation, while fading at the unexpected time points. Previewing our results, we did not find evidence that rhythmic temporal expectations elicit feature-specific prestimulus templates. We found instead a clear modality-specific (yet stimulus-non-specific) oscillating representation in the neural signals, demonstrating entrained rhythmic pre-activation of task-relevant sensory cortices. This sensory entrainment persisted after the offset of rhythmic stimulation. Our results demonstrate the existence of rhythmic nonspecific sensory pre-activation in the brain, highlighting the multitude of ways in which expectations can modulate neural activity.

## Materials and Methods

### Data and script availability

All data, as well as all presentation and analysis scripts, will be made freely available online upon publication, at the Donders Repository.

### Participants

Forty-two adult volunteers (16 male, average 27 years) participated in the experiment. Volunteers were excluded when they had more than 20% of no response trials (n=7) or had a dental wire (n=1). Thirty-four were included for the behavioural and MEG analyses. This sample size was determined a priori to ensure 80% power to detect a within-participant effect of medium size (d ≥ 0.5, paired t-test). This effect size should not be interpreted as an expected effect size, but we instead decided on this particular effect size as the minimum value we wanted to be sensitive for. All participants had normal or corrected-to-normal visual acuity, normal hearing, and health conditions consistent with the experiment. This study was approved under the general ethics approval (“Imaging Human Cognition”, CMO 2014/288) by CMO Arnhem-Nijmegen, Radboud University Medical Centre. All participants provided written informed consent.

### Apparatus

Computational routines were generated in MATLAB (The MathWorks) and stimuli were presented using “Psychtoolbox” (Brainard, 1997). A PROpixx projector (VPixx Technologies, Saint-Bruno, QC Canada) was used to project the visual stimuli on the screen, with a resolution of 1920×1080 and a refresh rate of 120 Hz, and the audio stimuli were presented through MEG-compatible ear tubes. Behavioural responses were collected via a MEG-compatible response box.

MEG was recorded from a whole-head MEG system with 275 axial gradiometers (VSM/CTF Systems, Coquitlam, BC, Canada) in a magnetically shielded room and digitized at 1200 Hz. Eye position data was recorded during the experiment using an Eyelink 1000 eye tracker (EyeLink, SR Research Ltd., Mississauga, Ontario, Canada) for further eye blink and saccade artefact rejection. During the session, head position was recorded and monitored online (Stolk, Todorovic, Schoffelen, & Oostenveld, 2013) by coils placed at the nasion, left and right ear. At the end of each block, participants were asked to reposition the head in case they moved more than 5 mm away from the initial position. MEG analyses were performed using FieldTrip software (Oostenveld, Fries, Maris, & Schoffelen, 2011) and repeated measures ANOVA were performed in JASP, Version 0.9.0 (JASP Team, 2018).

### Stimuli and general task

Participants performed auditory and visual discrimination tasks. In visual trials, the target was a grating of 3 degrees of visual angle, with a spatial frequency of 2 cycles per degree (cpd), random phase, and with one of six possible orientations (15, 45, 75, 105, 135, 165 degrees) surrounded by a magenta circle and presented centrally for 100 ms. In auditory trials, one of six possible pure tones (501, 661, 871, 1148, 1514, 1995 Hz) was presented for 100 ms as a target.

The target was always followed by a delay period, after which a probe stimulus was presented for 100 ms. The probe was similar to the target with the exception of the pitch/orientation feature. Participants had to judge whether the probe was tilted clockwise (CW) or counterclockwise (CCW) relative to the target in visual trials or whether the probe had a frequency higher or lower than the target in auditory trials. They always responded with a button press of either the index (lower/CCW) or middle finger (higher/CW) of their right hand.

Stimulus presentation timing was either non-rhythmic or rhythmic, in order to manipulate temporal expectations. The experimental session started with the non-rhythmic trials and the rhythmic trials were presented subsequently.

### Non-rhythmic trials

The non-rhythmic trials began with a central white fixation point (0.4 degrees of visual angle) with a surrounding cyan circle (3 degrees), and after a random inter-trial interval (ITI) chosen from a uniform distribution between 0.6 s and 1.6 s, a target was presented for 100 ms. Three seconds after target onset, the probe was presented, and participants had to indicate their response. There was no time limit for the responses. Participants received performance feedback for 400 ms and a new ITI started immediately (Figure 1 A). There were 24 trials in each block. Six blocks were randomly presented to participants (three auditory and three visual), resulting in a total of 72 trials per attended sensory modality. To ensure participants understood the task, they performed at least six easy practice trials for each condition before the procedure. Practice trials were not included in analysis.

**Figure 1:**
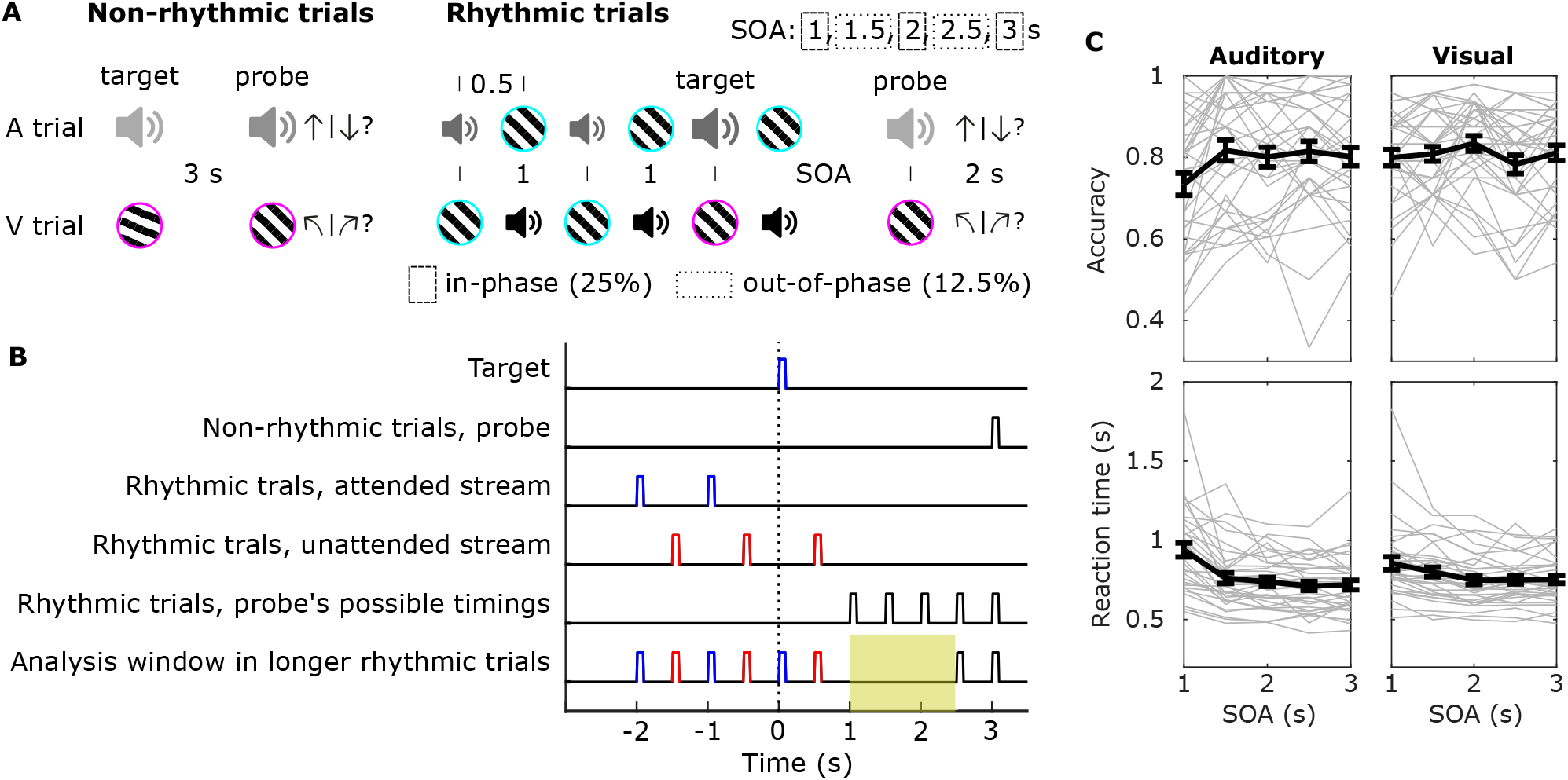
Experimental design and behavioural results. A) Schematic of the non-rhythmic and rhythmic trials. In both tasks, there were visual and auditory blocks. In visual blocks (V trial), participants had to discriminate whether the probe had a counterclockwise or clockwise tilt compared to the target. In auditory blocks (A trials), they had to judge whether the pitch was lower or higher. The non-rhythmic trials had a fixed configuration of one target followed by one probe. In rhythmic trials, the relevant stimulus was presented several times (3 to 6) with a fixed interval between presentations (1s) in order to induce a 1Hz entrainment. Interleaved and irrelevant to the task, a second stream of stimuli in the unattended sensory modality stimuli were presented. The last relevant stimulus (target) of the sequence was marked by a change in an irrelevant feature to warn participants about the oncoming probe presentation. The warning signal was a higher volume sound (illustrated by a larger sound icon, A trials) or a magenta outline (V trials). The interval between target and probe could be 1, 1.5, 2, 2.5 or 3 s with the respective probabilities: 25%, 12.5%, 25%, 12.5%, 25%. B) Timeline of the stimuli in the trials and the delay in the main analysis window. The yellow shadow represents the analysis window (delay from 1 to 2.5 s). C) Accuracy and reaction times (mean and standard error of the mean in bold) for the rhythmic task, as a function of SOA and attended modality. Individual participant data are shown in lighter grey.

Performance in non-rhythmic trials was also used to calibrate visual and auditory parameters for the following rhythmic experimental manipulation. The staircase method was QUEST as implemented in the Palamedes Toolbox (Prins & Kingdom, 2018), which we set to achieve a performance around 75%. For both conditions, a Cumulative Normal function with a lapse rate of 0.1 and the mean of the posterior were used as the staircase parameters. For the visual condition, the beta value(slope) was 1 and the prior alpha (threshold) range was a normal distribution with mean of 10 degrees, standard deviation of 5, ranging from 0 to 20 degrees. During the experiment, the chosen alpha value plus a random value from a normal distribution function (mean 0, std 1) was added or subtracted from the target orientation value. For the auditory condition, the beta value was 100, and the prior alpha range was a normal distribution with mean of 0.1, standard deviation of 0.1, ranging from 0 to 0.2. During the experiment, the chosen alpha value plus a random value from a normal distribution function (mean 0, std 0.01) were multiplied with the target pitch value and the resulting value was added or subtracted from the target value. Beta and alpha prior values were based on prior piloting results.

### Rhythmic trials

In rhythmic blocks, each trial started with a fixation point and a cyan circular border (Figure 1 A). Half a second later, the first attended-modality stimulus was presented for 0.1 s. This stimulus was presented several times (3 to 6, balanced and randomly chosen per trial) with a fixed interval between them (1 s) to create a rhythmic stream. Interleaved and irrelevant to the task, a second stream of stimuli in the other sensory modality stimuli was presented (unattended stream). Therefore, the interval between the onset of adjacent stimuli was 0.5 s. The last attended stimulus of the sequence (target) was marked by a change in an irrelevant feature to warn participants that the next presented stimulus would be the probe. The design with repeated stimuli was important to induce the rhythmic expectations. We used a variable number of repetitions in order to induce an uncertainty in the exact time point of target presentation, thereby making the rhythm itself (rather than one particular time point) all the more relevant to attend. The warning signal was a higher volume sound (auditory trials, equal to non-rhythmic target volume) or a magenta outline (visual trials, instead of cyan outline). Half a second after target onset (400 ms after the target offset), an unattended modality stimulus was always presented for 100 ms. Probes had a positive or negative difference in orientation or pitch based on the output from the previous staircase procedure and were also marked by the magenta outline or by the volume increase. The interval between target and probe (SOA) could be 1, 1.5, 2, 2.5 or 3 s with the respective probabilities: 25%, 12.5%, 25%, 12.5%, 25%. Participants were informed at the beginning of the experimental session that the timing of the probe was most likely to follow the attended rhythm, i.e., it would likely to occur in phase with it. A new trial with a new modality-relevant stimulus would appear 2 s after the probe. Each block consisted of 12 trials.

There were 16 auditory and 16 visual rhythmic blocks. There were 192 trials for each sensory modality. Volunteers performed 48 trials for each in-phase delays (1,2,3 s) and 24 trials for each anti-phase SOAs (1.5, 2.5 s) in each sensory modality condition. The blocks were always presented in a pseudo-random order, where no more than 3 same type blocks could be presented in a row, and participants were instructed about the block-type before its beginning.

### Behavioural analysis

Trials where participants did not respond within 3 s post-probe range were treated as incorrect trials in accuracy analyses and were excluded from reaction time (RT) analyses. Accuracy scores were arcsin-transformed before all statistical tests to improve normality. Both measures were submitted to a 2×5 repeated measures ANOVA with modality (auditory or visual) and the five SOAs as factors. Mauchly’s test of sphericity was performed, and Greenhouse–Geisser correction was applied in case of sphericity violation. Holm correction for multiple comparisons was performed for all post-hoc analyses, when applicable.

### MEG pre-processing

An anti-aliasing low-pass filter at 600 Hz was used during the online MEG recordings. Non-rhythmic trials were segmented between 0.2 s before the target until 0.5 s after the probe. Rhythmic trials were segmented between 0.2 s before the first stream stimulus until 0.5 s after the probe. After segmentation, synthetic 3rd order gradient correction was applied and the channel- and trial-wise mean was subtracted from the traces. Trials with eye movements, muscular activity and with an unusually high variance were excluded from the further analyses using a semi-automatic procedure (rejected trials: mean = 7.2%, SD = 3.3%). Sensors showing an unusually high variance were rejected following the same procedure (rejected sensors: mean = 2.8%, SD = 1.2%). After artifact rejection, data were off-line downsampled from 1200 Hz to 400 Hz to speed up analyses, followed by an independent component analysis to identify and remove residual eye, heart and other muscular components. A discrete Fourier transform was used to suppress line noise at 50 Hz and its harmonics, 100 Hz and 150 Hz.

### Planar combined event related fields

For the analysis of event-related fields (Figure 2), all trials were low pass filtered at 35 Hz and baseline corrected from −0.1 to 0 s. All non-rhythmic trials (number of visual trials ranged from 54 to 72, median = 69; number of auditory trials ranged from 65 to 72, median = 71) were considered from −0.2 s to 2.5. To better illustrate the activity during the delay of the rhythmic trials, only 2.5 and 3 SOAs trials (number of visual trials ranged from 52 to 72, median = 64.5; number of auditory trials ranged from 60 to 72, median = 68) were considered from −2.2 s to 2.5 s, including the 3 repetitions of attended and unattended stimuli in a trial. We chose to include only longer SOA trials for the evoked-field analysis to have a sufficiently long “clean" window (i.e., a period without any stimulation). For each participant, trials were time-lock averaged. MEG axial gradiometers were then transformed to planar configuration (Bastiaansen & Knösche, 2000) and combined as the root-mean-square of horizontal and vertical sensors. The combined planar activity from participants were averaged in the end.

**Figure 2:**
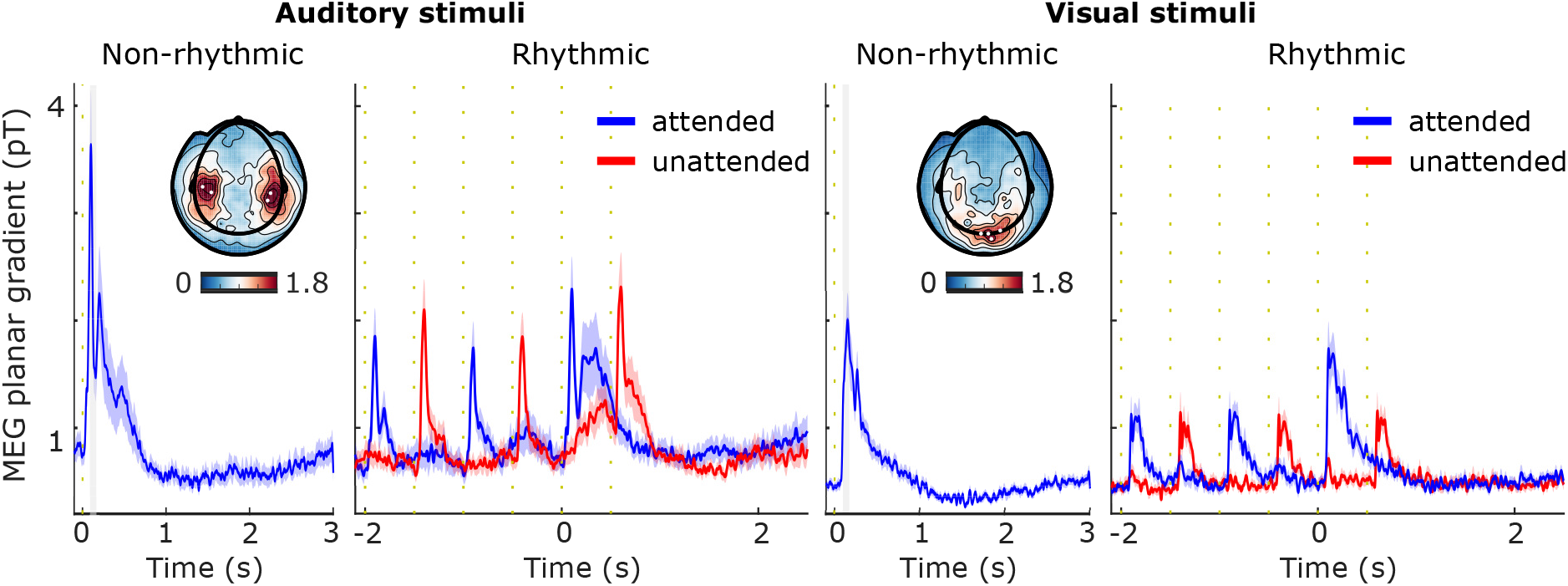
Event related field (mean and standard error of the mean) of non-rhythmic, attended (blue) and unattended (red) rhythmic longest-SOA trials in auditory and visual sensors. Time 0 represents the target presentation. Inset topographies illustrate the average activity related to non-rhythmic (auditory and visual) targets from 100 to 200 ms (lighter grey box period), and the white dots represent the most active sensors within this time window. The corresponding sensors were selected for representing the ERFs. Vertical dashed line (yellow) indicates a stimulus occurrence.

### Multivariate pattern analyses

MVPA were performed using linear discriminant analysis (LDA) as implemented in MVPA-Light toolbox (https://github.com/treder/MVPA-Light). Features consisted of activity in the MEG sensors (267 ± 3 sensors). Feature scaling was performed as pre-processing step in all analyses: data were normalized for each time point, across trials, using z-score transformation based only on the training set. Final scores were calculated based on the distances estimated by LDA from the six classes’ centroids in multiclass classification or from the hyperplane in the two-class classification.

### Specific feature classification: Temporal generalisation

We were interested in investigating the temporal evolution of potential feature-based preparatory activity that could occur during the delay, and in whether its generation was related to the temporal expectations. We looked at this first through the lense of temporal generalisation (i.e., train on various time points during stimulus period, test on various time points during delay), which would allow us to detect an (pre-/re-)activation of any part of the stimulus-evoked pattern during the delay period. Only the rhythmic trials with the longest delay periods (2.5 s and 3 s) were used in the testing set here, as well as all non-rhythmic trials (3 s delay). Testing trials were locked to the target and were segmented from −.1 s to 2.5 s in rhythmic trials and from −.1 s to 3 s in non-rhythmic trials. The training set was the presentation time window (-.1 to .5 s) from all stimuli presented in the rhythmic shortest trials (SOA 1, 1.5, 2 s). Segments were baseline corrected based on pre-target window (-.1 to 0 s). Given that six classes of orientations and six classes of tones were presented, we fitted two multiclass LDA: an orientation classification model (visual) and a tone classification model (auditory).

For non-rhythmic trials, orientation classification model was tested in visual trials and the tone classification model was tested in auditory trials. At the end, the average of the decoding scores between auditory and visual trials was computed. For rhythmic trials, both sensory models were tested in each trial since all rhythmic trials contained one visual and one auditory presented feature. Depending on the test trial task, the auditory and visual scores were assigned an attended or unattended label. For example, in an orientation discrimination (attend-visual) trial, the grating to be decoded was the visual attended feature and the tone was the auditory unattended feature, while in a pitch task (attend-auditory) trial the labels were attended auditory and unattended visual. At the end, scores from visual and auditory trials were averaged in relation to their rhythmic attention labels.

### Specific feature classification: Temporal decoding

To potentially increase sensitivity to stimulus-specific activity during the delay period, we also performed a decoding analysis while training on those time points of maximal stimulus-evoked activity. Trials were locked to the target and segmented until the probe moment. Subsampling by averaging 32.5 ms temporal windows (13 points in time) was applied to improve the signal-to-noise ratio. We performed classification in a leave-one-trial-out cross-validation approach. Accordingly, excluding the rhythmic test trial, all stimuli segments from rhythmic and non-rhythmic trials were used for training. Segments were baseline corrected based on pre-target window (-.1 to 0 s). The activity used in the training set was the average activity of each axial sensor from 0.1 to 0.2 s after a stimulus presentation. This training time period was chosen based on a previous study showing that the visual template effect (Kok et al., 2017) resembles the ERF peak activity (Figure 3 B). Depending on the test trial task, the auditory and visual scores were assigned an attended or unattended label. Trial length was SOA-condition dependent.

**Figure 3:**
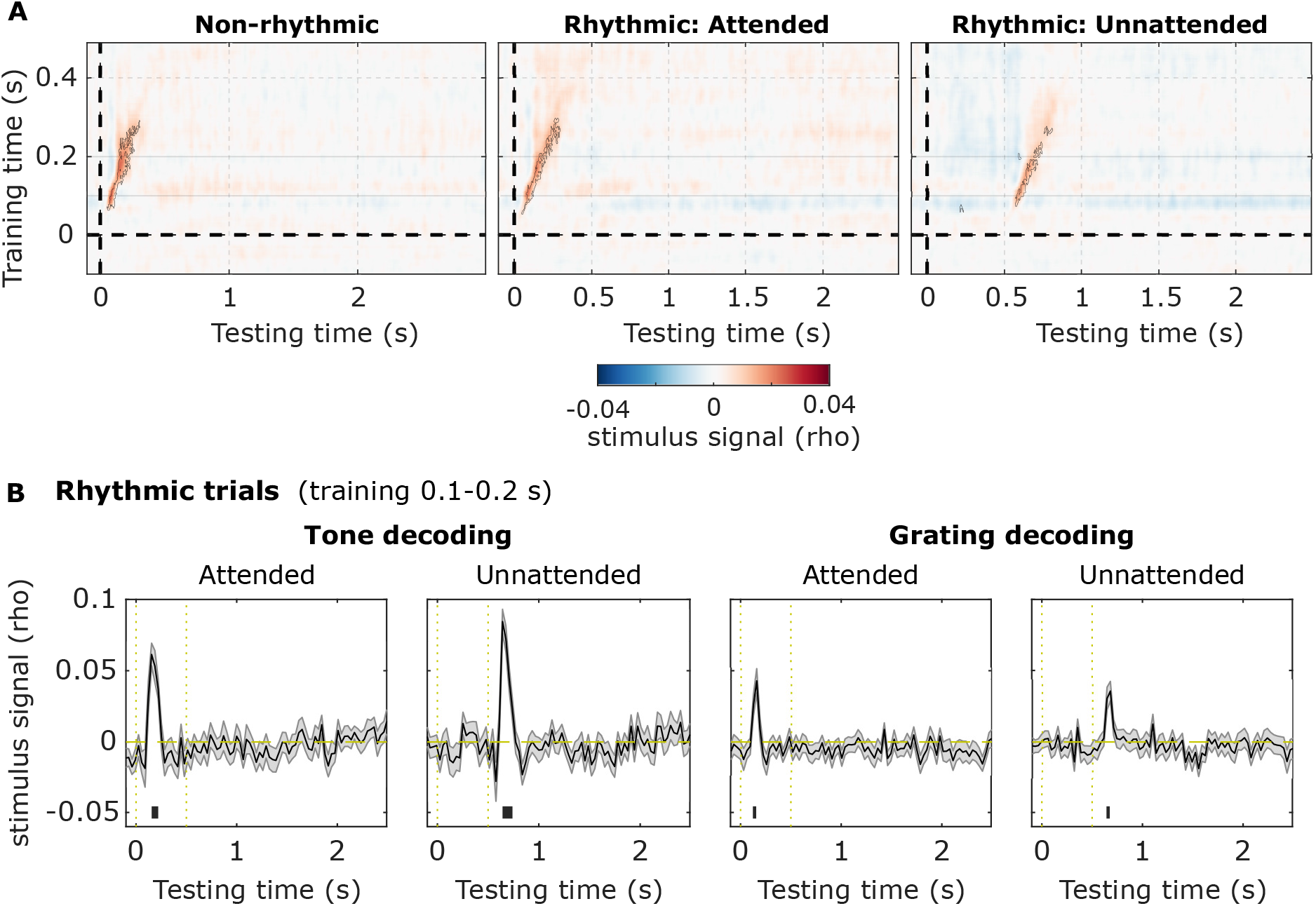
Multivariate decoding of stimulus-specific information (pitch/orientation). A) Temporal generalisation matrices for non-rhythmic, attended and unattended longest-SOA rhythmic trials. Target presentation occurred at 0 s, and the unattended stimulus (rhythmic trials) at 0.5 s. Significant clusters (p <0.05) are contoured in black, thus illustrating a momentary transient feature-specific signal after stimulus presentation. B) Leave-one-trial-out cross-validation results using the averaged sensor activation from 0.1 to 0.2 s as training data. Vertical dashed line (yellow) indicates attended (t = 0 s) and unattended (t = 0.5 s) stimulus occurrence and the black bars indicate the significant clusters (all p <0.01). Only actual stimulus periods, and not the stimulus-absent delay period, had scores higher than chance.

### Specific feature classification: Score

The scores were the estimated rho from a Spearman rank correlation test between the estimated distances and an “ideal distances matrix” (Auksztulewicz, Myers, Schnupp, & Nobre, 2019). This ideal matrix was the expected trial distance, or rank, for each of the six classes’ centroids. For the visual condition, with orientation being a circular variable, the expected distance rank was the lowest, 0, for the correct label (i.e. 15 degrees), 1 to the two closest label neighbours (165 and 45 degrees), 2 for the middle far two classes (135 and 75 degrees) and 3 for the further class (105 degrees). The auditory condition matrix was different from the visual given that frequency is a linear variable. The lowest distances were drawn along the diagonal and gradually higher ones for further off-diagonal positions.

### Sensory modality nonspecific classification

The existence of a more general, but sensory-specific, preparatory activity could be analysed during the delay as well. Since we were mainly interested in the temporal evolution of the modality-specific signals during this post-stimulus period, only longest-delay trials (2.5 s and 3 s) for rhythmic and non-rhythmic conditions were used in this analysis. They were target locked, cut between −0.2 s until the probe moment, and baseline corrected based on pre-target window (-.1 to 0 s). Here we used a two-class LDA, again with a temporal generalisation approach. To keep computational time manageable, data was downsampled to 200 Hz before this analysis. The scores here were the distances to the decision hyperplane calculated by LDA.

### Statistics

Scores from trials were averaged within each time point for each participant. To assess significant differences from chance, we used cluster-based permutation tests based on paired t-scores (Maris & Oostenveld, 2007), with 1000 random permutations, channel neighbours as defined in the FieldTrip neighbourhood template for the CTF275 machine (no additional threshold for minimum number of neighbouring channels (minnbchan = 0), cluster alpha of 0.01 and cluster statistics as the maximum of the summation.

## Results

We investigated the role of temporal expectations in a multisensory task. In different blocks, participants (n = 34) had to perform either a pitch (Auditory blocks) or orientation (Visual blocks) discrimination task. The first part of the experimental session consisted of simple discrimination trials, where participants were presented with a single visual or auditory target (target) followed by a unisensory probe of the same modality (Figure 1 A). Participants had to judge whether the probe was tilted clockwise or anti-clockwise relative to the target in visual trials or whether the probe had a frequency higher or lower than the target in auditory trials. We refer to these trials as the “non-rhythmic” trials. During these non-rhythmic trials, difficulty was adjusted according to an adaptive staircase procedure (Watson & Pelli, 1983), to titrate the difference in grating angle and tone frequency to an appropriate difficulty level (75%) for the rest of the experiment. Participants’ thresholds in the visual task ranged from 5.49° to 19.98° tilt (Q1/4 = 9.82°, Q2/4 = 12.33°, Q3/4 = 18.75°) and in the auditory task ranged from 0.38% to 19.93% pitch difference (Q1/4 = 3.01%, Q2/4 = 14.93%, Q3/4 = 18.14%) which were smaller values than the steps used for the visual (30° of orientation) and auditory targets (min of 24% of the target pitch).

In the second part of the experimental session, probes were preceded by a stream of 2 Hz alternating visual and auditory stimuli (rhythmic trials). In a blockwise fashion, participants had to either pay attention to the visual stream (1 Hz) and perform the visual orientation task or pay attention to the auditory stream (1 Hz) and perform the pitch discrimination task. Visual and auditory stimuli presented in the stream had the same orientation and pitch. They could have one of six possible orientations and one of six possible pitches.

The last stimulus in the attended stream was the target (again of the same orientation/pitch as the preceding stream) and was identifiable to the participant by either a coloured ring (visual) or increased volume (auditory). Critically, probes could appear after one of five possible stimulus onset asynchrony (SOA) intervals related to the target: 1, 1.5, 2, 2.5, or 3 s (Figure 1 A and B). Integer intervals were in-phase relative to the attended stream, while 1.5 and 2.5 s were in anti-phase. The probability of presentation at the in-phase SOAs was 25%, whereas it was 12.5% at the anti-phase SOAs, making it more likely that the probe would be presented in-phase with the attended stream. With this design, including a clear delay period (Figure 1 B), we could test whether specific (i.e. decodable orientation and tone signals) and/or non-specific sensory (visual or auditory evoked activity without decodable orientation/tone signals) activation continued after the stimulation period. We hypothesised that feature-based expectations could be generated during the delay, since targets roughly predicted the probes’ features (i.e., the probe’s orientation/pitch would be very similar to the target and preceding stream). As a result, they might prompt sensory templates during both the non-rhythmic and rhythmic conditions, while we expect these templates, if present, to be rhythmically modulated only in the rhythmic condition. Furthermore, temporal expectations might be stronger in the rhythmic trials, and, more importantly, qualitatively different, since it is known that single-interval and rhythmic temporal predictions rely on distinct neural mechanisms (Breska & Ivry, 2018). For these reasons, we investigated the presence and possible rhythmic modulation of sensory patterns during these conditions.

### Behavioural performance

We first tested whether, in the present task, rhythmic presentation of targets resulted in a rhythmic modulation of perceptual performance. We measured performance based on accuracy and reaction time (RT) (Figure 1 C). Accuracy was lowest for the earliest SOA (mean = 76.7%, SEM = 1.8%), an effect most pronounced for the attend-auditory blocks. This was backed up by a significant main effect of SOA (F(4,132) = 6.37, p = 6.172e-04, *ω*^2^ = 0.03), as well as an interaction of SOA and attended modality (F(4,132) = 5.02, p = 8.451e-04, *ω*^2^ = 0.02). Overall accuracy was not different between the modalities (main effect of sensory modality: F(1,33) = 0.02, p = 0.886, *ω*^2^ = 0), and despite the interaction, SOA affected accuracy in both the attend-visual and attend-auditory blocks (simple main effects analysis of SOA, auditory: F(1) = 7.45, p = 1.92e-05; visual: F(1) = 2.89, p = 0.025). However, only the first SOA differed from the other intervals (1.5 s: 81.3 ± 1.8%, t(33) = −3.39, p = 0.015, d = −0.582; 2 s: 81.8 ± 1.8%, t(33) = −4.76, p = 3.711e-04, d = −0.82; 3 s: 80.7 ± 1.8%, t(33) = −3.93, p = 0.004, d = −0.67), except from 2.5 s (79.9 ± 1.9%, t(33) = −2.34; p = 0.181, d = −0.4). Additionally, Bayes factor analysis shows that the data were about four times more likely under the null than under the alternative hypothesis (BF10 = 0.246, Bayesian t-test) when computing in-phase (2 and 3 s) and anti-phase (1.5 and 2.5 s) accuracy scores from the four remaining SOAs.

Reaction times decreased with increasing SOA (1 s: 896 ± 38 ms; 1.5 s: 780 ± 28.5 ms; 2 s: 743 ± 24.5 ms; 2.5 s: 730 ± 23.5 ms; 3 s: 735 ± 23.6 ms; main effect of SOA F(4,132) = 32.27, p = 2.234e-09, *ω*^2^ = 0.12). Reaction times were not significantly different between attended sensory modalities (F(1,33) = 0.09, p = 0.763, *ω*^2^ = 0). We did observe an interaction of SOA and attended modality (F(4,132) = 7.57, p = 0.001, *ω*^2^ = 0.02), while SOA affected reaction time in both the attend-visual and attend-auditory blocks (simple main effects analysis of SOA, auditory: F(1) = 28.98, p = 2.725e-17; visual: F(1) = 12.36, p = 1.438e-08). Responses for the shortest SOA (1 s) were slower than for the other SOAs (post-hoc t-tests, 6.05<t(33)<7.32, 2.14e-7<p<5.779e-6, 1.04<d<1.26), and responses for the 1.5 s SOA were slower than those for the longer SOAs (2 s: t(33) = 2.82, p = 0.032, d = 0.48; 2.5 s: t(33) = 3.29, p = 0.014, d = 0.56; 3 s: t(33) = 3.18, p = 0.016, d = 0.55), while response times for the SOAs >1.5 s did not differ among one another (−0.59<t(33)<1.31, 0.6<p<0.88, −0.1<d<0.22). When computing in-phase (2 and 3 s) and anti-phase (1.5 and 2.5 s) reaction times from the four last SOAs, there was no evidence for any hypothesis (BF10 = 1.833, Bayesian t-test). Taken together, behavioural performance provides no evidence for a significant rhythmic modulation of perceptual performance, but instead points toward a hazard rate effect.

### Stimulus-specific information is decodable from MEG sensors during stimulation only

Next, we turned our attention to the neural consequences of interleaved multisensory rhythmic stimulation. Figure 2 shows the event-related fields for MEG sensors approximately overlying auditory and visual cortices, in all different conditions. As expected, auditory and visual stimuli elicited pronounced event-related fields at the auditory and visual associated sensors. The evoked activity returned to baseline levels approximately 1 s after stimulus presentation.

To quantify whether the neural signals contained stimulus-specific information (i.e., information about which of the six auditory pitches or visual orientations was present), we performed a multivariate pattern analysis. Specifically, we trained classifiers on the period (−100 to 500 ms) from all stimuli presented in rhythmic trials with SOAs 1, 1.5, and 2 s, and quantified how well these generalised to both the stimulus and delay periods of the rhythmic trials with the longest SOAs (2.5 s and 3 s), as well as to the non-rhythmic trials. Train and test data here are thus fully independent. We investigated the cross-temporal generalisation of these signals between the full stimulus training period to the combined stimulus and delay testing period.

We observed a strong feature-specific signal when training and testing the classifier on similar time points (Figure 3 A). In all conditions, there were high levels of stimulus information in the diagonal (non-rhythmic: from 55 ms to 305 ms post-stimulus, p <0.001; attended rhythmic: from 50 ms to 295 ms post-stimulus p <0.001; unattended rhythmic: from 75 ms to 275 ms post-stimulus, p = 0.002; p-values estimated using cluster-based permutation tests). However, we found no evidence of a generalisation of this activity to other time points in the delay period or in anticipation of events.

This first analysis suggested that sensory representations elicited by a specific feature did not generalise to the delay period. We found a momentary and transient feature-specific signal that peaked after stimulus presentation. To test whether increasing the size of the training set might increase our sensitivity to a potentially missed result, we repeated the classification procedure within the rhythmic conditions only. In this new analysis: (1) data from all trials (excluding a single trial) were used as a training set (i.e. we used a leave-one-trial-out procedure); (2) we used as the training time the period around the ERF peak activity (100 to 200 ms). Similar to our previous analysis, we found that only periods around stimulus presentation had scores higher than chance (auditory attended: from 155 ms to 220 ms, p <0.001; auditory unattended: 143 ms to 240 ms, p <0.001; visual attended: from 123 ms to 155 ms, p = 0.008; visual unattended: from 143 ms to 175 ms, p = 0.004; Figure 3 B). We did not find feature-specific sensory activation during the delay period and patterns evoked by specific orientations and tones were restricted to periods of stimulus-driven activity.

### Sensory cortices pre-activate rhythmically during delay periods

It is known that rhythmic stimulation can entrain neural activity in related sensory areas. Having found no evidence that this entrainment is feature-specific, we next explored whether multimodal rhythmic stimulation induced non-specific, yet modality-specific, rhythmic activation of sensory cortices during the delay. We again used a temporal generalisation approach, this time to decode the attended modality (visual or auditory). The modality decoding reflects how strongly a pattern corresponding to visual activation is present in the sensors versus how strongly a pattern corresponding to auditory activation is present considering the attentional label as the direction of evidence. A significant positive decoding means that neural sources typically processing information of the attended sensory modality are more active than neural sources typically processing information of the unattended sensory modality (in reverse for significant negative decoding scores).

The attended sensory modality was significantly decodable from the signal across several time points in both rhythmic and non-rhythmic trials (cluster-based permutation tests: non-rhythmic p <0.001; rhythmic p <0.001, Figure 4 A). Importantly, the modality signal extended throughout the delay periods in both types of trials (Figure 4 A). Its temporal evolution differed between conditions: the modality signal showed clear rhythmicity only in the rhythmic trials, indicating a pre-activation signal related to temporal expectation that was not immediately driven by any stimulus. The negative values during early training periods in rhythmic trials can be explained by noting that the unattended (i.e., different modality) stimulus was presented at those times.

**Figure 4:**
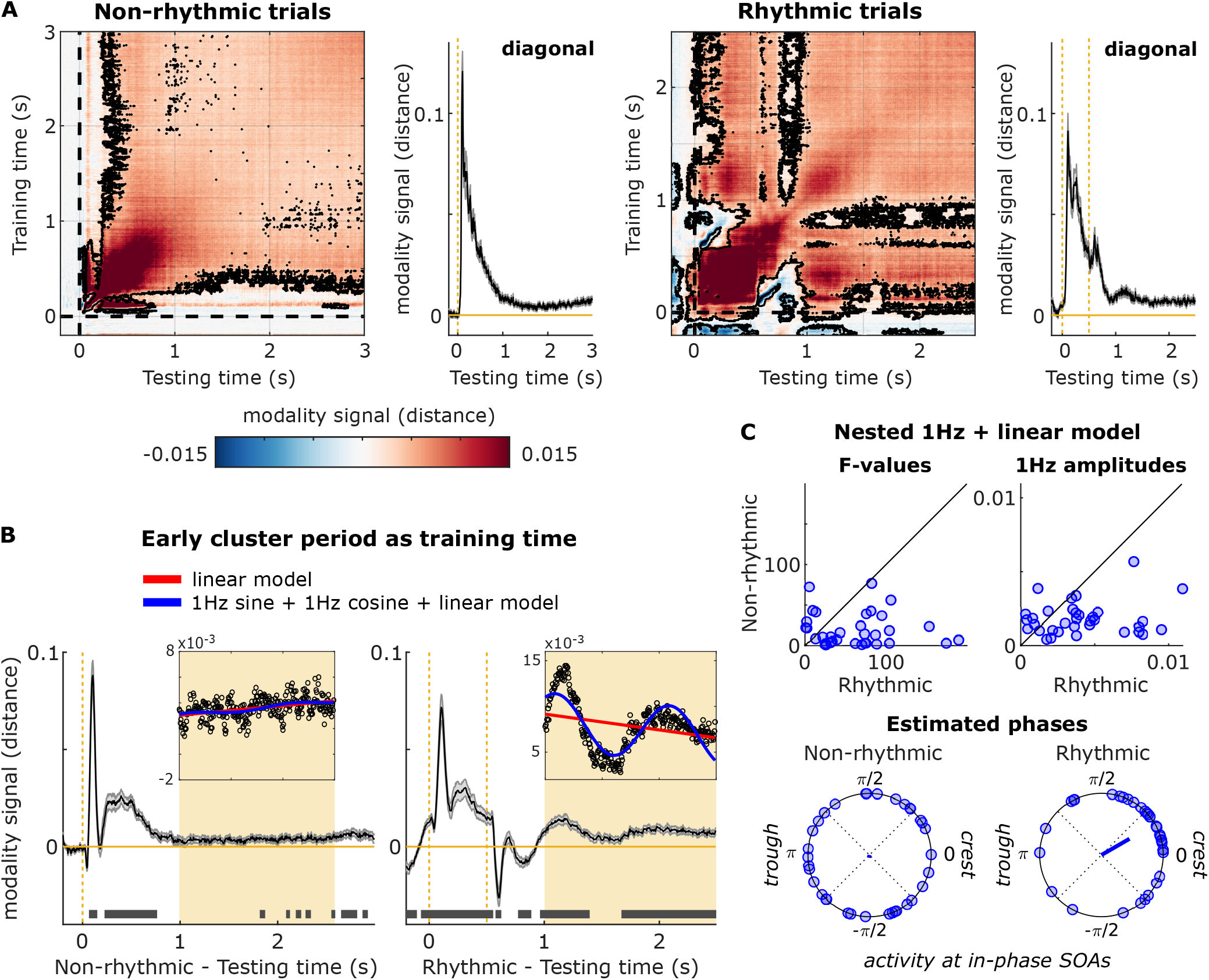
Multivariate decoding of relevant modality information (visual/auditory). A) Temporal generalisation matrices indicating a generalised sustained activation (significant clusters surrounded by a black line). In non-rhythmic trials, a significant cluster (training time 0.08 to 0.13 s) illustrates that an early sensory representation pops out at the end of the interval. B) Temporal evolution of such early sensory representation in both non-rhythmic and rhythmic conditions. Significant clusters are indicated as grey bars. In rhythmic trials, modality signal oscillated after stimulation period. Two models were fitted into the no-stimulus delay data (1 s to 2.5 s, inset) for non and rhythmic conditions. C) F-stats of nested models and the amplitude of the fitted sinusoid. This indicates that 1Hz oscillation model explains better the modality signal behaviour in rhythmic than in non-rhythmic trials. Furthermore, there is no phase preference in non-rhythmic trials, but phases are highly clustered in rhythmic trials, indicating a better representation at in-phase/highly expected delays.

In both types of trials, we observed an early pattern of activity (training time 0.08 s to 0.13 s) that was strong and generalised to different testing times throughout the delay period (Figure 4 A). To study the temporal dynamics of this activity in more detail, we further analysed performance over time for a classifier trained on this time window, in both types of trials. Figure 4 B shows how modality activity evolves. We observed a clear oscillatory modulation of decoding scores, which can be seen during the delay period for rhythmic trials, and which was absent for non-rhythmic trials. Critically, the last stimulus presented in this period was at 0.5 s, with no other stimulation after that. Additionally, 400 ms after the offset of the unattended stimulus (at 1 s), the modality signal reverses to the attended modality by staying largely positive and significant during the delay. This suggests that such modality signal during the delay reflects preparatory sensory activity rather than the previous stimulus evoked activity.

An oscillatory modulation of the modality signal was clearly present in the grand-averaged data (Figure 4 B). We next assessed whether this rhythmicity was reliably present across participants, by fitting two models to the activity in the delay period (1 to 2.5 s) for each participant. The first was a linear model with intercept and slope as free parameters. The second model was a combination of a linear function with a 1 Hz sine and 1 Hz cosine function (which is equivalent to a 1 Hz sinusoid with phase as a free parameter) (Zoefel, Davis, Valente, & Riecke, 2019). The combined 1Hz-linear model provided a significantly better fit of the data for 26 out of 34 participants in the non-rhythmic condition (Wald test controlling for the extra degree of freedom; F values range: 0.88 to 76.72, critical F(2,295): 3.03), and for 31 out of 34 in the rhythmic condition (F values range: 0.61 to 189.47, critical F(2,295): 3.03). As the critical test of whether stimulus periodicity induced a rhythmic modulation of sensory cortex activation during delay periods, we compared the improvement in model fit that resulted from adding the sinusoid term between the rhythmic and non-rhythmic trials. The improvement of adding an oscillatory function was considerably higher for rhythmic than non-rhythmic trials (Wilcoxon signed-rank test of relative F-values across participants; Z = 3.77, p = 1.634e-04; Figure 4 C, top left). Furthermore, the model fits for the rhythmic trials had significantly higher 1 Hz amplitudes than those for non-rhythmic (Wilcoxon signed-rank test of 1Hz amplitudes across participants; Z = 3.86, p = 1.156e-04; Figure 4 C, top right).

If the delay-period oscillatory modality signal is the result of entrainment by the rhythmic stimuli, one would expect the phases of this signal to be consistent across participants, specifically for the rhythmic (and not the non-rhythmic) condition. This is indeed what we observed: phases were not significantly different from uniform in non-rhythmic trials (Rayleigh test; Z(33) = 0.07, p = 0.93), but we observed a clear phase concentration in rhythmic trials (average phase = 0.525 rad, Z(33) = 9.37, p = 4.783e-05; Figure 4 C, bottom panels). To rule out that the evoked activity of the irrelevant stimulus in the rhythmic condition at 0.5 s could be an explanation for a stronger phase-locking compared to the non-rhythmic, we also tested the phase consistency at later latencies. Phases were still consistent (Z(33) = 8.708, p = 1.016e-04) in the rhythmic condition when we considered the analysis window from 1.5 to 2.5 s (1 s after the last presented stimulus), while there was no phase consistency in the non-rhythmic condition, already in the window from 1 to 2 s (so also 1 s after the last presented stimulus; Z(33) = 0.5, p = 0.611).

Taken together, these results demonstrate that the rhythmic stimulation resulted in a rhythmic pre-activation of sensory cortices, which was consistent across participants. Importantly, this pre-activation was observed during the delay period, i.e. without any ongoing sensory stimulation, suggesting a true entrainment of endogenous neural signals.

## Discussion

In the present study, we investigated whether rhythmic temporal prediction interacts with feature-based expectations to induce rhythmic sensory templates for anticipated stimuli. Behaviourally, we found that temporal expectations improved performance, but not in a rhythmic manner. Using multivariate pattern analysis of feature-specific signals, we found that stimulus information was present only during stimulation and not during the delay period, contrary to our expectations. Instead, we observed feature-unspecific but modality-specific activity during the delay, reflecting a rhythmic pre-activation of the relevant sensory cortices that peaked at the expected, behaviourally relevant, moments.

Contrary to what we expected, performance was not modulated in line with the rhythm of the task. Although participants exhibited worse performance for the first SOA (both in response times and in accuracy), performance was not different between the other SOAs. According to Dynamic Attending Theory (DAT) (M. R. Jones & Boltz, 1989; M. R. Jones, Moynihan, MacKenzie, & Puente, 2002), in-phase intervals should lead to faster and more accurate responses than anti-phase intervals. In this study, we only found a general increase in performance as a function of delay. This is a well-known result called the variable foreperiod effect that can be explained by the increasing conditional probability of target occurrence with increasing SOAs, also known as the “hazard function” (Näätänen, 1970; Nobre et al., 2007; Nobre, 2010). One could argue that the absence of behavioural effect could be that the entrained rhythm was at 2 Hz by considering both streams and not at 1 Hz. Although possible, we do not have any evidence that supports this hypothesis from our neurophysiological data.

Several studies have found evidence in support of the DAT: performance is improved in rhythmic compared to arrhythmic conditions (Rohenkohl et al., 2012; Morillon, Schroeder, Wyart, & Arnal, 2016), and the phase of entrained neural oscillations by an external rhythm influences auditory (Henry & Obleser, 2012; Bauer, Bleichner, Jaeger, Thorne, & Debener, 2018) and visual perception (Cravo et al., 2013; Chota & VanRullen, 2019). Thus, there is a large literature suggesting that environmental rhythms can entrain attentional (i.e., endogenous, neural) rhythms and modulate perception (Henry & Herrmann, 2014). Nevertheless, results are not as clear when analysing post-entrainment effects, i.e., after the offset of the rhythm. Different studies have shown behavioural impairments (Hickok, Farahbod, & Saberi, 2015; Spaak et al., 2014), benefits (M. R. Jones et al., 2002; Barnes & Jones, 2000) or effects that were highly participant-dependent (Bauer, Jaeger, Thorne, Bendixen, & Debener, 2015; A. Jones, 2019) for in-phase versus anti-phase time points. Similar to our design, a recent study showed no rhythmic behavioural facilitation for orientation discrimination tasks (Lin et al., 2021). Differences in the task (detection/discrimination), the sensory modality (time/auditory/visual), and the stimulated frequency range (alpha/delta) could have led to this variety of different effects. Together with our null result regarding rhythmicity in post-entrainment behaviour, these results highlight the necessity for additional studies to understand the factors that determine the influence of rhythms on behaviour.

Previous studies have shown that feature-based expectations about an event can induce anticipatory activation templates in sensory cortex (Kok et al., 2014, 2017). Here we tested this possibility in different modalities (vision and audition), conditions (rhythmic and non-rhythmic), and levels of task relevance (attended or unattended). In all conditions, we found a similar pattern: stimulus-specific information could be decoded during the stimulation period only, and not during the following delay period. There are several differences between our and previous experiments, which might explain this discrepancy. One important difference may be the information to be stored. In previous studies, stimulus-specific pre-activation was found after an informative cue presented in anticipation of the target stimulus. Our task, in contrast, required the maintenance of target information (tone frequency or grating orientation) that needed to be later compared to a probe; thus, participants had already seen the target itself before the period of interest. Consequently, working-memory processes were involved during the delay period as well as potential anticipatory processes by considering the prospective nature of a memory (Nobre & Stokes, 2019). Classifiers were always trained on the stimulus evoked activity, since we were trying to detect anticipatory activity similar to stimulation, as in previous work. It has been argued that such stimulus-identical delay activity is not strictly necessary for working memory maintenance, and that information might have been stored in a different, possibly silent, format (Wolff, Ding, Myers, & Stokes, 2015; Mongillo, Barak, & Tsodyks, 2008; Stokes, 2015). It is possible that sensory templates may be instrumental for automatic associations between two events, while our paradigm favours a more prospective, silent, coding scheme. This would also fit with the behavioural task: unlike previous works, in our task, participants had to compare upcoming stimulation with what came before, thus an exact stimulus-specific pre-activation of the earlier stimulus might even impair behavioural performance. In line with this interpretation, it has been reported that neural reactivation increases serial biases (Barbosa et al., 2020), which would impair performance here. Finally, a possible reason for not finding stimulus-specific templates is that MEG might simply not be sensitive to pick these up. Since we found neither auditory nor visual templates, we believe this is unlikely, however, since visual orientation is well-known to be detectable in the MEG signal (Stokes, Wolff, & Spaak, 2015; Cichy, Ramirez, & Pantazis, 2015).

Although we did not find stimulus-specific anticipatory information, there was a clear pre-activation of the relevant sensory cortices. The early modality-specific signal was decodable in a rhythmic fashion in rhythmic trials, and it was also present close to the end of the delay period in non-rhythmic trials. Thus, the pre-activation was locked to the temporal structure of the task and peaked at the time points of expected stimulation. Such higher visual/auditory activity did not represent the stored orientation/tone information for the memory task, but it was likely generated by prospective memory processes related to probe’s anticipation. This modality-specific pre-activation is consistent with recent results that showed that even task-irrelevant information was better decoded when presented at moments close to a highly likely target presentation (Auksztulewicz et al., 2019). Analogous to these previous results, this boosting of early sensory modality representations in the MEG signal during the delay could be explained by increases in baseline excitability of task-relevant sensory areas (as opposed to task-irrelevant ones). Thus, the modality-specific decoding reflects a relative measure of neuronal excitability between auditory and visual cortex. It has previously been shown that endogenous neural oscillations in visual cortex bias perception through rhythmic fluctuations in baseline excitability (Iemi, Chaumon, Crouzet, & Busch, 2017). Our results suggest that similar fluctuations can be leveraged in a cross-modal setting to optimally prepare the brain for upcoming stimuli, specifically of a task-relevant modality. It should be noted that we cannot conclude with certainty that the rhythmic modulation of visual/auditory cortex excitability is due either to the visual, or the auditory stream, since the two streams were tightly phase-locked in our experiment. We believe the most likely interpretation to be a relative waxing and waning of excitability between the two modalities.

Previous studies have shown that neuronal population excitability states can be entrained to external rhythms as a preparatory mechanism for optimally processing upcoming stimuli (Lakatos et al., 2008; Schroeder & Lakatos, 2009; Henry & Obleser, 2012; Lakatos et al., 2013; Herrmann, Henry, Haegens, & Obleser, 2016), which is, in turn, under strong top-down control (Lakatos et al., 2019). It is important to distinguish entrainment from other factors such as superimposed evoked responses, resonance, and endogenous predictions (Guevara Erra, Perez Velazquez, & Rosenblum, 2017; Helfrich, Breska, & Knight, 2019), which may interact with entrainment (Haegens, 2020). In the present study, we used a decoding analysis of the expected sensory modality as a slightly different than usual approach to evaluate entrainment. We analysed the engagement of early sensory cortices during a silence period after two conditions: a single evoked stimulus, and a 1 Hz stream. Instead of computing the traditional Fourier transform to evaluate oscillatory power and phase consistency in neural data, we computed the relative neuronal excitability between visual and auditory cortex. It is a relative measure because the decoding score of the modality-signal corresponds to the amount of evidence for the attended sensory activation topography compared to the unattended one. We investigated whether rhythmic excitability shifts at 1 Hz were enhanced at post-entrained compared to post single stimulus periods. Our results are in line with previously published work in monkeys (Lakatos et al., 2008), which we extended by showing that: (1) the oscillatory pattern is present in the absence of external stimulation, and (2) the oscillatory pattern more strongly arises after a rhythmic stream than after a single stimulus. These two points, combined, strongly suggest that our results were not due to superposition of responses or a simple resonance mechanism. Although the combined sinusoid-linear model was also a better fit than the purely linear model for non-rhythmic trials, this improvement was considerably stronger in rhythmic trials. Furthermore, phase was scattered uniformly for the non-rhythmic trials, but consistent in rhythmic trials. Taken together, we conclude that the modality decoding signal on non-rhythmic trials did not contain a rhythmicity.

Lastly, in our experimental setup, in-phase moments were also moments in which there was a higher probability of target presentation. For this reason, it is not possible to dissociate the effects of locally stimulus-driven oscillatory entrainment from globally generated predictive signals introduced by the probability manipulation. Given that we observed a (non-rhythmic) increase of decoding scores towards the end of the delay period in non-rhythmic trials, globally generated predictions might explain part of our results. Whether these endogenous predictions are enhanced in the presence of rhythms, and/or whether they interact with local sensory oscillatory entrainment is still an open question that should be addressed in future studies.

In summary, our results add to the body of evidence showing that the brain extracts temporal regularities from the environment to optimally prepare in time for upcoming stimuli. Importantly, we demonstrate that one specific mechanism for such temporal attunement in a visual/auditory cross-modal setting is the phasic modulation of excitability in early visual and auditory cortex, in lockstep with the environment. We furthermore show that the occurrence of stimulus-specific, actively maintained, early sensory anticipatory templates, as reported previously, appears to depend on the specifics of the task at hand, and is not a universal phenomenon.

## Acknowledgments

This work was supported by São Paulo Research Foundation (FAPESP, grants 2016/04258-0 and 2018/08844-7 awarded to LCB, and grant 2017/25161-8 awarded to AMC), by The Netherlands Organisation for Scientific Research (NWO Veni grant 016.Veni.198.065 awarded to ES and Vidi grant 452-13-016 awarded to FPdL), and by the European Research Council (ERC), under the European Union’s Horizon 2020 research and innovation programme (grant agreement No. 678286 to FPdL). The funders had no role in study design, data collection and analysis, decision to publish or preparation of the manuscript.

## Author contributions statement

All authors conceived the experiment. LCB performed the experiments and LCB and ES analysed the data. All the authors wrote and reviewed the manuscript.

